# Phenotypic and genotypic characterization of antibiotic Resistant Gram Negative Bacteria isolated in Tabuk City, Saudi Arabia

**DOI:** 10.1101/2021.02.01.429288

**Authors:** Tarig M.S. Alnour, Elmutuz H. Elssaig, Eltayib H. Ahmed-Abakur, Faisel M. Abuduhier, Khalid A. S. Alfifi, Mohammad S. Abusuliman, Tawfiq Albalawi

## Abstract

Antimicrobial surveillance and identifying the genetic basis of antimicrobial resistance provide important information to optimize patient care. The present study was analytical cross sectional study aimed to determine the prevalence of MDR, XDR, PDR and extended-spectrum β-lactamases genes (SHV, CTX-M and TEM) among Gram-negative bacteria isolated in Tabuk, Saudi Arabia. A total number of 386 non-duplicate Gram-negative isolate were collected. Identification and susceptibility testing were done using automation system (BD Phoenix™). The extracted DNA were subjected to multiplex polymerase chain reaction (PCR). The results showed that only 15 (3.9%) of isolates were fully susceptible, the overall prevalence of XDR, MDR, PDR was 129 (33.4%), 113 (29.3%) and 48(12.4%) respectively. High resistant rate was observed against the antibiotic agents of cephalosporins class 79.3% followed by the agents of penicillins class 69.4%. The most dominant gene was bla SHV which detected in 106/386 (27.5%) isolates followed by bla CTX-M 90/386 (23.3%). Bla CTX-M showed significant relation with all used antibiotic except ampicillin/clavulanic acid, aztreonam, cefoxtin, and meropene. The isolates which showed most frequent resistant genes were *Klebsiella pneumoniae 90/124 (72.6%), A. baumanni* 37/67 (55.2%), *and P.mirabilis* 24/44 (54.5%). These findings underscores the need for optimization of current therapies and prevention of the spread of these organisms.

## 1. Introduction

Antimicrobial resistance is described as a condition in which the pathogen escape from the stress of the antibiotic exposure (1). The increasing incidence of antimicrobial resistance is the key concern globally and considered main obstacle in the treatment of patients suffering from bacterial infections.(1, 2) It has been estimated that about 1.3 to 2 fold rise in mortality caused by antimicrobial resistant bacteria compared to susceptible infections. (1).

A dramatic evolution has occurred in the significance of infections caused by Gram-negative bacteria (GNB) and associated with considerable mortality (1, 2, 3). The efficiency of the current prophylactic and empiric antibiotic treatment is compromised by the emergence of pan drug resistant (PDR), extensively drug resistant (XDR) and multidrug resistant (MDR) Gram-negative bacteria (GNB) (2, 3). The ability of escaping from the antimicrobial effects may be contributed to the nature of these organisms, which are heterogeneous, complex group of plasmids-borne and rapidly evolving enzymes, which are capable of hydrolyzing cephalosporins, penicillins, aztreonam and monobactams (4).

The American society of infectious diseases identified six top priority dangerous pathogens producing extended spectrum β-lactamases (ESBLs). Three of these six pathogens are antibiotic resistant Gram-negative bacteria; *Pseudomonas aeruginosa*, *Acinetobacter baumannii* and Enterobacteriaceae (5). ESBLs have been classified into three major groups; bla SHV, bla CTX-M and bla TEM (2), Ting et al., 2013 stated that bla TEM, bla SHV and bla CTX-M genes are super-resistant extended spectrum β-lactamases (6). In 2017, the WHO published list of global priority pathogens, a catalog of twelve species of bacteria grouped according to their antibiotic resistance under three priority tiers; critical, high, and medium. The critical group involved three pathogens that were Gram-negative bacilli; *Acinetobacter baumannii*, *Pseudomonas aeruginosa*, and *Enterobacteriaceae*, (7). The WHO also notified that the level of resistance to antimicrobial drugs used to treat common infections is reaching a crisis point. If world administrations do not control infections in order to slow down the growth of drug resistance, entire populations could be wiped out by superbugs (8, 9). However, the availability of regional information on the resistance rate is fundamental to implementing efficient treatment protocols against infectious pathogens and may help to prevent infections with multidrug resistance pathogens at the local level (10, 11). Therefore, the present study was aimed to determine pattern of antimicrobial resistance and to detect ESBL genes among Gram-negative bacteria isolated in Tabuk city, Saudi Arabia.

## 2. Materials and Methods

The present study was analytical cross sectional study, conducted in King Fahad Specialist Hospital and prince Fahad Bin Sultan Research Chair (University of Tabuk), Saudi Arabia. A total number of 386 non-duplicate Gram-negative isolates were collected in order to determine the prevalence of MDR, XDR, PDR and to detect extended-spectrum β-lactamases genes bla SHV, bla CTX-M and bla TEM.

### Identification and susceptibility test

Depending on the origin of the samples, each sample cultured on suitable medium/ media from: MacConkey agar, CLED agar blood agar, Chocolate agar or Brain Heart infusion broth. Then they were incubated aerobically at 37°C for 24 to 48 hours except for the blood culture, which was incubated for 5 to 7 days in broth medium. Growth of corresponding organisms were further sub-cultured for purification purpose. The significant growth was identified to the species level. Identification and susceptibility testing were done using automation system (BD Phoenix™). Identified strains were tested in vitro against several antimicrobial classes including carboxypenicillin (Ticarcillin/Clavulanic acid), Penicillinase resistant penicillin (Ampicillin/Sulbactam and Piperacillin/Tazobactam), Cephalosporins (Ceftazidime and Cefepime), Aztreonam, Carbapenems (Ertapenem, Imipenem and Meropenem), Aminoglycoside (Amikacin, Gentamicin and Tobramycin), Fluoroquinolones (Ciprofloxacin and levofloxacin), Minocycline (Tetracyclin), Glycylcycline (Tigecycline), polymyxin E (Colistin), and sulpha drugs (Trimethoprim/Sulfamethoxazole). Antimicrobial selection for testing depends on types of isolates and site of samples which done automatically by the program. Based on susceptibility test of the above mentioned antibiotic classes the isolates were characterized as MDR, XDR and PDR.

Moreover thirteen agents of five antimicrobial classes which represented the most commonly used antibiotics; carboxypenicillin, penicillinase resistant penicillin, cephalosporins, aztreonam and carbapenems were subjected for further study in order to determine the relation between these antibiotics and ESBL genes, these agents included: ampicillin, ampicillin/clavulanic acid, ticarcillin/clavulanic acid, aztreonam, piperacillin/tazobactam, cefalotin, cefoxitin, ceftazidime, ceftrixone, cefepime, imipenem, meropenem and ertapenem. Quality control and maintenance were achieved according to the manufacturer’s guidelines.

The BD Phoenix™ automated identification and susceptibility testing system empowers workflow efficiency using automated nephelometry, which results in a standardized isolate inoculum and a reduction in potential technologist error along with accurate, reliable and rapid detection of known and emerging antimicrobial resistance (12).

### Detection of Antimicrobial Resistance Genes

DNA was extracted from whole 386 GNB isolates using boiling technique; few colonies from each isolate were mixed with molecular biology-grade water (Eppendorf, Hamburg, Germany), the mixture were centrifuged at 15,000 × *g* for 5 min. The supernatant was discharged and the pellet was re-suspended in molecular biology-grade water (Eppendorf, Hamburg, Germany) and subjected to boiling at 100°C in a water bath for 20 min, then cooled and centrifuged at 15,000 × *g* for 60s before it was stored at −20°C.

Multiplex polymerase chain reaction (PCR) were done to determine the presence of three ESBLs genes encoding bla TEM, bla SHV, bla CTX-M. Each extracted DNA was tested against the three sets of specific primers in a single test (Table 1). Amplification was performed in a final volume of 25 μL containing 5μl of template DNA, 0.5 μl Taq polymerase, 1.0 μl of each primers and 0.2 μl dNTP mixture (10 mM), and finally the volume was completed to 25 μL by molecular biology-grade to reach volume of 25 μl. The PCR protocol was run according to the following protocol; essential denaturation at 95°C for 5 minutes followed by 40 cycle of denaturation at 95°C for 30s, annealing at 60°C for 30s and extension at 72°C for 1 minutes, the final step was extension at 72°C for 5 minutes. PCR product was run on 2% ethidium bromide agarose gel electrophoresis, and examined with gel imaging system, bands pattern was observed and interpreted according to their size (Table 1).

**Table 1:**
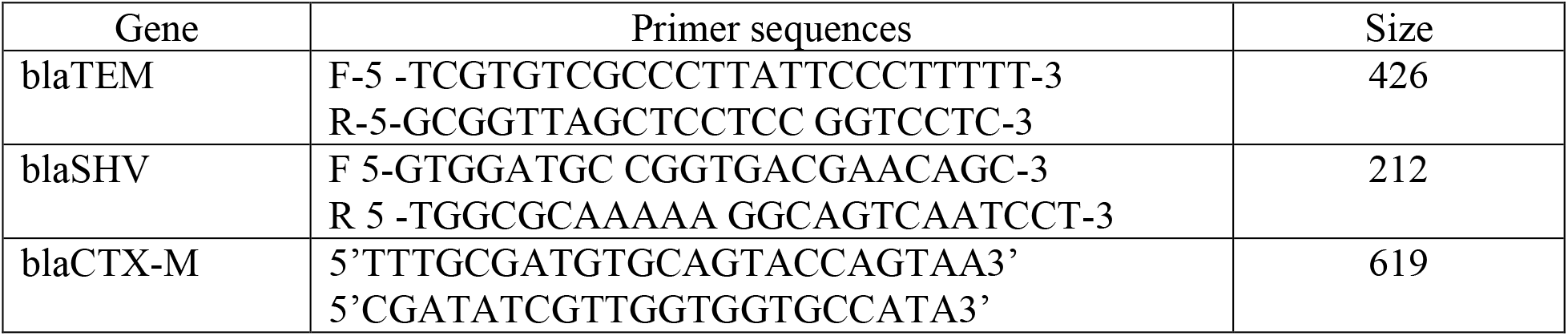
the primers sequences of ESBLs genes and corresponding size

#### Analysis

The proportion of resistant for each antibiotic was calculated as the sum of resistant antibiotic relative to the sum of susceptible and resistant. The proportion of resistant class of antimicrobial was represent the mean of resistance of all antimicrobial agents belong to that class. Chi-square tests was performed to determine the relation between ESBLs genes and antibiotic resistant using SPSS version 22. *P* value <0.05 was considered significant.

The isolates which showed susceptibility to all groups of antimicrobial agents were classified as susceptible, while those showed resistant to one group were classified as mono-resistant. The isolates resistant to one drugs in two groups were classified as resistant to two antimicrobial group. Multidrug resistant (MDR) denoted when the isolates shown resistance to 3 or more antimicrobial group but susceptible to 2 or more. The US Centers for Disease Control and Prevention (CDC) and European Centre for Disease Prevention and Control (ECDC) have defined bacteria as pan drug resistant (PDR) when they are non-susceptible to all agents in all antimicrobial categories and as extensively drug-resistant (XDR) when they are non-susceptible to at least one agent in all but two or fewer antimicrobial categories (13).

The ethical clearance for this study (UT-86-10-2019) was obtained from research ethics committee, University of Tabuk (Saudi Arabia).

## Result

Table no 2 showed the frequency of isolates along with patterns of occurrence of resistant ESBLs genes; the results revealed that the most common isolates were *Klebsiella pneumoniae* 124(32.1%), followed by *A. baumanni* 67(17.4%), *E.coli* 51(13.2%), *P.aeruginosa* 50(13.0%) and *P.mirabilis* 44(11.2%).

**Table 2:**
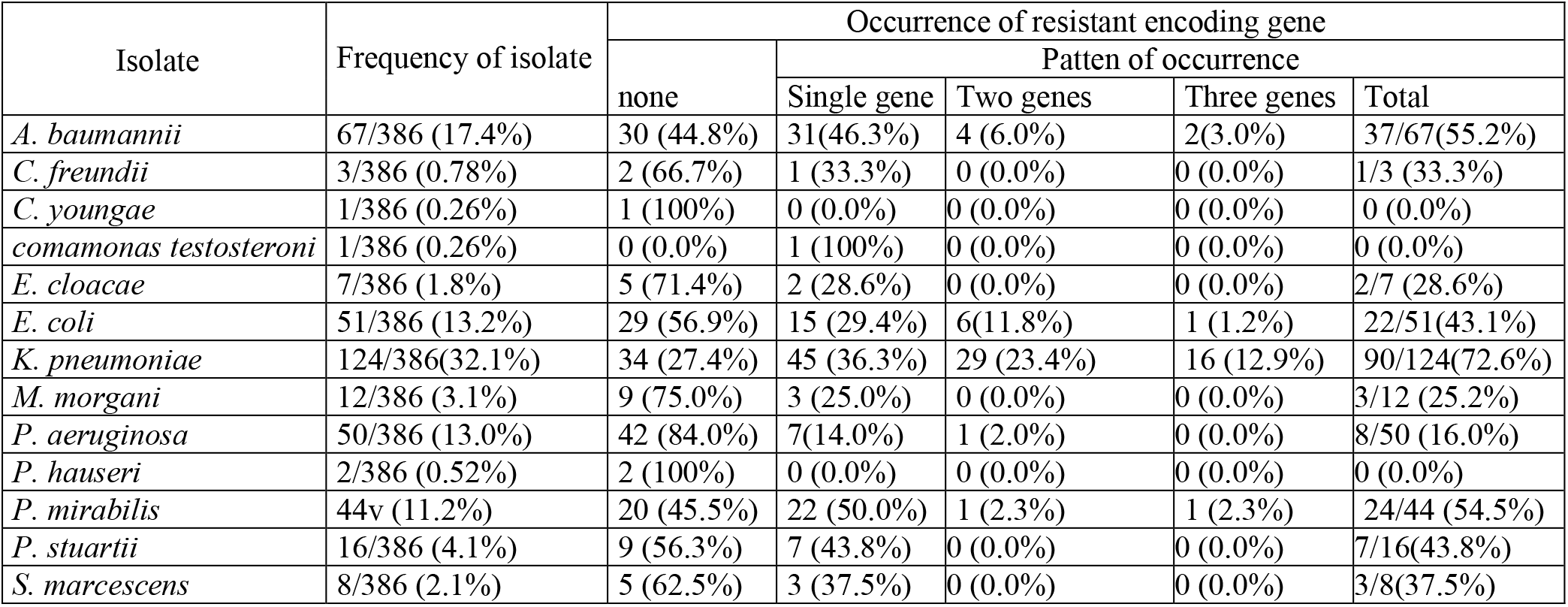
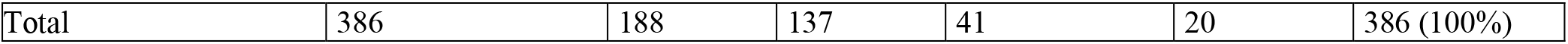
Frequency of isolates and patterns of occurrence of resistant ESBLs genes

| Isolate | Frequency of isolate | Occurrence of resistant encoding gene |  |  |  |  |
| --- | --- | --- | --- | --- | --- | --- |
|  |  | none | Patten of occurrence |  |  |  |
|  |  |  | Single gene | Two genes | Three genes | Total |
| *A. baumannii* | 67/386 (17.4%) | 30 (44.8%) | 31(46.3%) | 4 (6.0%) | 2(3.0%) | 37/67(55.2%) |
| *C. freundii* | 3/386 (0.78%) | 2 (66.7%) | 1 (33.3%) | 0 (0.0%) | 0 (0.0%) | 1/3 (33.3%) |
| *C. youngae* | 1/386 (0.26%) | 1 (100%) | 0 (0.0%) | 0 (0.0%) | 0 (0.0%) | 0 (0.0%) |
| *comamonas testosteroni* | 1/386 (0.26%) | 0 (0.0%) | 1 (100%) | 0 (0.0%) | 0 (0.0%) | 0 (0.0%) |
| *E. cloacae* | 7/386 (1.8%) | 5 (71.4%) | 2 (28.6%) | 0 (0.0%) | 0 (0.0%) | 2/7 (28.6%) |
| *E. coli* | 51/386 (13.2%) | 29 (56.9%) | 15 (29.4%) | 6(11.8%) | 1 (1.2%) | 22/51(43.1%) |
| *K. pneumoniae* | 124/386(32.1%) | 34 (27.4%) | 45 (36.3%) | 29 (23.4%) | 16 (12.9%) | 90/124(72.6%) |
| *M. morgani* | 12/386 (3.1%) | 9 (75.0%) | 3 (25.0%) | 0 (0.0%) | 0 (0.0%) | 3/12 (25.2%) |
| *P. aeruginosa* | 50/386 (13.0%) | 42 (84.0%) | 7(14.0%) | 1 (2.0%) | 0 (0.0%) | 8/50 (16.0%) |
| *P. hauseri* | 2/386 (0.52%) | 2 (100%) | 0 (0.0%) | 0 (0.0%) | 0 (0.0%) | 0 (0.0%) |
| *P. mirabilis* | 44v (11.2%) | 20 (45.5%) | 22 (50.0%) | 1 (2.3%) | 1 (2.3%) | 24/44 (54.5%) |
| *P. stuartii* | 16/386 (4.1%) | 9 (56.3%) | 7 (43.8%) | 0 (0.0%) | 0 (0.0%) | 7/16(43.8%) |
| *S. marcescens* | 8/386 (2.1%) | 5 (62.5%) | 3 (37.5%) | 0 (0.0%) | 0 (0.0%) | 3/8(37.5%) |
| Total | 386 | 188 | 137 | 41 | 20 | 386 (100%) |

The isolates showed high resistant rate against the antimicrobial agents of cephalosporins group 79.3% followed by the agents of penicillins 69.4% while against the agents of carbapenems they exhibited 32.5 % resistance rate. In term of individual antimicrobial agent; the isolated Gram-negative bacteria revealed a high resistance rate against ampicillin 262(93.2%) followed by aztreonam 87(90.6%) and cefalotin 157(90.2%). Only 15 (3.9%) of isolates were fully susceptible to all used antimicrobials. The overall prevalence of MDR, XDR, PDR was 113 (29.3%), 129 (33.4%) and 48(12.4%) respectively (Table no 3).

**Table 3:** Phenotypic characterization of antibiotic resistance among Gram-negative bacteria

| Phenotypic characterization of isolate | Frequency | Percentage |
| --- | --- | --- |
| Mono-resistant | 28 | 7.2 |
| Resist to 2 antimicrobial group | 53 | 13.7 |
| MDR | 113 | 29.3 |
| XDR | 129 | 33.4 |
| PDR | 48 | 12.4 |
| Fully Sensitive | 15 | 3.9 |
| Total | 386 | 100% |

Screening for resistance genes showed that, most of Gram-negative isolates harbored the resistance genes 198/386(51.3%) while isolates free from resistant genes were 188 (48.7%) (Table 2). Bla SHV was most dominant gene which detected in 106/386 (27.5%) isolates followed by bla CTX-M and bla TEM which detected in 90/386 (23.3%) and 78/386 (20.2%) isolates, respectively. Single resistant gene was detected in 137/386 (35.5%) isolates, coexistence of two genes were detected in 41/386 (10.6%) isolates while triple genes were present in 20 (5.2%) isolates. Bla CTX-M showed significant relation with all used antibiotic except ampicillin/clavulanic acid, aztreonam and cefoxtin while bla SHV exhibited significant statistic relationship with Piperacillin/ Tazobactam, Cefalotin, ceftazidime, cefepime, imipenem and meropenem. Bla TEM displayed significant relation only to ampicillin/clavulanic acid, ceftazidime, cefepime and imipenem. Ceftazidime and cefepime agents of cephalosporins class were least effective agents as they showed significant relation to the three resistant genes (Table 4). The isolates which showed the most frequent resistant genes were *K. pneumoniae* 90/124 (72.6%), *A. baumanni* 37/67 (55.2%), *E. coli* 22/51 (43.1%), *P. aeruginosa* 8/42(19.0%) and *P. mirabilis* 24/44 (54.5%) (Table 2).

**Table 4:** Phenotypic resistance along with frequency of ESBLs genes

| Beta lactam groups | Antimicrobials | Resistance Nr (%) | Genes |  |  |
| --- | --- | --- | --- | --- | --- |
|  |  |  | blaSHV<br>Nr (%) | blaTEM<br>Nr (%) | blaCTX-M<br>Nr (%) |
| Penicillinase resistant penicillins and carboxypenicillin | Ampicillin | 262/282 (93.2) | 92/262 (35.1) | 60/262 (22.9) | 78/262 (29.8)<br>$P = 0.005$ |
| | Ampicillin/ clavulanic acid | 78/141 (55.3) | 46/78 (58.9) | 21/78 (26.9)<br>$P = 0.022$ | 32/78 (41.0) |
| | Ticarcillin/Clavulanic acid | 96/161 (59.6) | 24/96 (25) | 18/96 (18.8) | 21/96 (21.9)<br>$P = 0.003$ |
| Aztreonam | Aztreonam | 87/96 (90.6) | 24/87 (27.6) | 23/87 (26.4) | 11/87 (12.6) |
| Monobactam / sulbactam | Piperacillin/ Tazobactam | 168/249 (67.5) | 59/168 (35.1)<br>$P = 0.006$ | 41/168 (24.4) | 58/168 (34.5)<br>$P = 0.000$ |
| Cephalosporins | Cefalotin | 157/174 (90.2) | 61/157 (38.9)<br>$P = 0.043$ | 37/157 (23.6) | 53/157 (33.8)<br>$P = 0.017$ |
|  | Cefoxitin | 88/174 (50.6) | 42/88 (47.7) | 22/88 (25) | 29/88 (33) |
| | Ceftazidime | 325/386 (84.2) | 98/325 (30.2)<br>$P = 0.016$ | 75/325 (23.1)<br>$P = 0.002$ | 87/325 (26.8)<br>$P = 0.002$ |
| | Ceftriaxone | 185/210 (88.1) | 64/185 (34.6) | 46/185 (24.9) | 64/185 (34.6)<br>$P = 0.021$ |
| | Cefepime | 322/386 (83.4) | 99/322 (30.7)<br>$P = 0.014$ | 75/322 (23.3)<br>$P = 0.001$ | 88/322 (27.3)<br>$P = 0.001$ |
| Carbapenems | Imipenem | 201/375 (53.6) | 56/201 (27.9)<br>$P = 0.003$ | 52/201 (25.9)<br>$P = 0.016$ | 51/201 (25.4)<br>$P = 0.020$ |
| | Meropenem | 158/381 (41.5) | 55/158 (34.8)<br>$P = 0.042$ | 33/158 (20.9) | 47/158 (29.7)<br>$P = 0.032$ |
|  | Ertapenem | 2/80 (2.5) | 0/2 (0) | 1/2 (50) | 0/2 (0) |

## Discussion

Bacteria resistant to different classes of antimicrobial agents are a main threat to humanity and high risk, which may return the world towards pre-antimicrobial era (13). While active surveillance systems are set up in many countries in Europe, USA and Asia, little is reported on antimicrobial resistance status among Gram-negative bacteria in the Middle East, Africa and Saudi Arabia (14). The present study determined the prevalence of MDR, XDR, PDR and extended-spectrum β-lactamases genes (TEM, SHV, CTX-M) among Gram-negative bacteria in Saudi Arabia.

The third generation cephalosporin such as ceftazidime and cefoperazone marked by stability to the common beta-lactamases of Gram-negative bacilli and these compounds are highly active against Enterobacteriaceae (15). Although the isolated Gram-negative bacilli in this study showed high resistant rate to the antibiotic agents of cephalosporins class 79.3% followed by the agents of penicillinase resistant penicillins which showed 69.4% resistant, while the agents of carbapenems had least resistant rate; the highest resistance rate was reported against ampicillin (93.2%) followed by aztreonam (90.6%) and cefalotin 157(90.2%). Similar trend of resistance was observed by <u>Ruppé</u> etail 2015 who owing the dramatic increase in the rates of resistance to third-generation cephalosporins to spread of plasmid-borne extended spectrum beta-lactamases (ESBLs) in *Enterobacteriaceae* and to occurrence of sequential chromosomal mutations, which may lead to the overproduction of intrinsic beta-lactamases, hyper-expression of efflux pumps, target modifications and permeability alterations in non-fermenting Gram-negative bacteria (16). The serious finding in our study was emerging of carbapenems resistant. Carbapenems agents considered the most active and potent agents against multidrug-resistant (MDR) (17), this finding is totally contradictory to that reported by Zaman et al., 2015 who determined the susceptibility pattern of Gram-negative bacilli isolated from a Teaching Hospital in Jeddah, Saudi Arabia and reported (100%) sensitive of enterobacteriaceae to carbapenems (18). However WHO have been listed carbapenem-resistant *Enterobacteriaceae* among the top tier of the antibiotic resistant that pose the greatest threat to human health (7).

The overall prevalence of MDR, XDR and PDR in this study was 29.3%, 33.4%, and 12.4% respectively. Several studies were conducted in Saudi Arabia and showed high resistance rate among Gram-negative bacteria, most of these studies focused on susceptibility per individual pathogen (1, 9, 11, 14, 19, 20) However, Ibrahim 2018, reported higher rate of MDR in Southwest Saudi Arabia (67.9%) (11). the present study showed high PDR and less XDR compared to that reported by Mohapatra et al 2018 (13). Increasing antimicrobial resistant in Saudi Arabia may also be owing to increased cross geographic transmission of drug resistant. Saudi Arabia is capital of Islamic world and has great number of expatriates, which makes it a potential center for the import and export of multi resistant strains (21), a recent study by Leangapichart et al (2016) showed that returned travelers from Hajj had acquired MDR *A. baumannii* and NDM producing *E. coli* during the Hajj event (22).

Our findings showed that more than half (51.3%) of isolated Gram-negative harbored with resistant genes while isolates free from resistant genes were 48.7%. Only 3.9% of isolates were fully susceptible to the used antimicrobials. This finding in alignment with Munita and Arias report (2016) (23) and with Patil et al (2019) reults, who showed increase in the resistant rate among gram-negative pathogen and stated that Gram-negative bacteria are continuously evolving mechanisms to deactivate clinically important antimicrobial drugs by acquisition of resistance elements such as bla SHV, bla TEM and bla CTX-M (2). However antimicrobial resistant is an outcome of multifaceted microbial interactions such as microbial characteristics to gain resistance genes, selective pressure owing to inappropriate use and widespread of antibiotics, resistance may arise by the acquisition of de-novo mutation during treatment or by acquisition of integrative or replicative mobile genetic elements that have evolved over time in microbes in the natural ecosystem (23).

Our results indicated that bla SHV was most prevalent resistant gene which detected in (27.5%) of isolates followed by bla CTX-M (23.3%). Similar results indicated bla CTX-M and bla SHV as the most prevalent genotypes of ESBLs producing gram-negative pathogens were reported in several countries (2, 15, 24). Asokan et al (2019), reported that the bla CTX-M gene indicating bacterial evolution due to cover prescription or weak enforcement of existing antibiotics policies (25). The present study showed that *Klebsiella pneumoniae* (72.6%), *A. baumanni* (55.2%), *P.mirabilis* (54.5%) *E.coli* (43.1%), and *P.aeruginosa* (19.0%), were the highest isolates harbored with of ESBLs genes, these findings were in alignment with Maina et al 2012 (15), Ibrahim 2018 (11) and Asokan et al (2019) (25). However Enterobacteriaceae such as *K. pneumoniae*, *E. coli*, *Proteus spp, P. aeruginosa*, were naturally competent and can uptake naked DNA from the environment in suitable conditions (2).

In our study bla CTX-M showed significant association to all used antibiotic agents of cephalosporins, this finding is in alignment with Maina et al (2102), who stated that CTX-M type extended-spectrum β-lactamases (ESBLs) showing resistance to third and fourth-generation cephalosporins and to aztreonam (15).

### Conclusion

infections caused by multi-resistant Gram-negative pathogens negatively influence patient outcomes and costs. This study showed that only 3.9% of isolates had susceptibility to all used antibiotics, high resistant rate was observed against the antimicrobial agents of cephalosporins class and penicillinase resistant penicillin. The most dominant gene was bla SHV.

## Acknowledgments

The authors would like to thank deanship of scientific research (DSR) University of Tabuk, Kingdom of Saudi Arabia for sponsoring the current research (project number S‑1440‑0339). We are very grateful for the Microbiology staff of King Fahad Specialist Hospital for their valuable efforts in sample collection. Thanks are also offered to Prince Fahad Bin Sultan Research Chair and Medical laboratory Technology staff, faculty of medical applied sciences, Tabuk University for their valuable support.

